# Molecular dynamics simulations of HIV-1 matrix-membrane interactions at different stages of viral maturation

**DOI:** 10.1101/2023.10.01.560363

**Authors:** Puja Banerjee, Kun Qu, John A. G. Briggs, Gregory A. Voth

## Abstract

Although the structural rearrangement of the membrane-bound matrix (MA) protein trimers upon HIV-1 maturation has been reported, the consequences of MA maturation on the MA-lipid interactions are not well understood. Long-timescale molecular dynamics (MD) simulations of the MA multimeric assemblies of immature and mature virus particles with our realistic asymmetric membrane model have explored MA-lipid interactions and lateral organization of lipids around MA complexes. The number of stable MA-PS and MA-PIP2 interactions at the trimeric interface of the mature MA complex is observed to be greater compared to that of the immature MA complex. Our simulations identified an alternative PIP2 binding site in the immature MA complex where the multivalent headgroup of a PIP2 lipid with a greater negative charge binds to multiple basic amino acid residues such as, ARG3 residues of both the MA monomers at the trimeric interface, and HBR residues (LYS29, LYS31) of one of the MA monomers. Our enhanced sampling simulations have explored the conformational space of phospholipids at different binding sites of the trimeric interface of MA complexes. Unlike the immature MA complex, the 2’ acyl tail of two PIP2 lipids at the trimeric interface of the mature MA complex is observed to sample stable binding pockets of MA consisting of helix4 residues. Together, our results provide molecular-level insights into the interactions of MA trimeric complexes with membrane and different lipid conformations at the specific binding sites of MA protein before and after viral maturation.

**Significance:** HIV-1 maturation facilitates the conversion of a newly formed immature virus particle to a mature infectious virion and initiates a new round of infection. The contributions of specific protein-lipid interactions in the HIV-1 assembly process are well recognized, however, the interactions of matrix protein lattice with the membrane before and after HIV-1 maturation are yet to be fully understood. Based on our simulated data, supported by prior experimental observations, the present study provides a molecular-level understanding of possible altered binding mode of PIP2 lipids after viral maturation. Identification of protein-lipid specific interactions, and lipid sorting data obtained from our long-time and large-scale atomistic MD simulations advance the understanding of the HIV-1 matrix and membrane maturation.

## Introduction

During the late phase of the Human Immunodeficiency Virus type 1 (HIV-1) replication cycle, the newly synthesized major structural protein, Gag, binds to and multimerizes on the inner leaflet of the host cell plasma membrane (PM), leading to viral assembly, budding, and release of immature virus particles(1). Subsequent to the budding process, newly formed immature virions undergo the proteolytic maturation process, resulting in the cleavage of Gag into its structural proteins (2,3). Gag protein is composed of several structural domains having different roles in viral replication and infectivity. N-terminal myristoylated matrix (Myr-MA) domain mediates membrane binding of Gag and envelope (Env) glycoprotein incorporation into virus particles (4–9). Past studies have indicated that the capsid (CA) domain plays a critical role in Gag oligomerization, whereas the nucleocapsid (NC) domain promotes higher-order multimerization of Gag through the binding to viral genomic RNA (gRNA) during the viral assembly process (10–14). The virion maturation process leads to a major structural rearrangement of all three folded domains and a dramatic change in viral morphology that alters the protein-protein, protein-lipid, and protein-RNA interactions (1,11,15–19). In the mature infectious virion, the matrix (MA) domain remains associated with the virion membrane, while capsid protein (CA) forms the outer shell of the core capsid structure, and the nucleocapsid (NC) domain condenses with the viral genomic RNA (gRNA). Furthermore, maturation of the matrix domain is believed to be associated with the rearrangement of the Env glycoprotein and membrane maturation (19,20).

It has been recognized that the HIV-1 membrane differs notably from the producer cell PM. Inner leaflets of the viral membrane are enriched in anionic phospholipids, e.g., phosphatidylinositol 4,5-bisphosphate [PI(4,5)P_2_ or PIP2], which enhances the membrane-binding affinity of the MA domain (21,22). MA-plasma membrane interactions are facilitated by several key factors (i) non-specific electrostatic interactions between basic amino acid residues of MA and anionic membrane lipids (phosphatidylinositol(PI), phosphatidylserine(PS)), (ii) specific interactions between MA protein and phosphatidylinositol-4,5-bisphosphate [PI(4,5)P_2_], and (iii) hydrophobic interactions with lipid tails mediated by the N-terminal Myr group of MA, a post-translational modification covalently attached to the N-terminal glycine residue (23–30). This Myr moiety can adopt two different conformations: sequestered in a hydrophobic pocket of MA and a Myr-exposed state (31). NMR (Nuclear Magnetic Resonance) experiments (9,29) and MD simulations (32) have revealed that MA-PIP2 interactions and MA trimerization facilitate Myr exposure. While the inner leaflet of the viral membrane forms microdomains of anionic phospholipids induced by the MA-lipid interactions, the outer leaflet forms raft-like lipid domains, enriched in sphingomyelin (SM) and cholesterol (33). Further, these nanodomains in the two leaflets of the asymmetric bilayer are proposed to be correlated by trans-bilayer coupling (34,35). The impact of the altered MA-MA interactions after viral maturation on the membrane lateral organization is yet to be explored.

In a prior cryo-electron tomography (cryo-ET) study of MA lattice organization in the immature and mature HIV-1 particles, a rearrangement of the MA trimer-trimer interface upon maturation was reported, and altered MA-PIP2 interactions were observed (19). In the immature virion, MA trimer-trimer interactions are governed by N-terminal residues, helix1, and 3_10_ helix residues. On the other hand, in the mature virion, the MA trimer-trimer interface is stabilized by the interactions between the basic residues of the HBR (highly basic region) loop with the acidic residues of the N-terminus of helix4 and the 3_10_ helices of adjacent MA monomers. Unlike the immature virion, the cryo-ET density map (EMDB ID: 13088) of the mature virus particle shows lipid density at the PIP2 binding pockets; however, cryo-ET data does not allow us to identify specific lipid molecules. In the past, experimental studies by Saad *et al.*, Shkriabai *et al.,* and Chukkapalli *et al.* demonstrated different MA-PIP2 binding modes (29,36,37). Among these, the extended PIP2 lipid configuration reported by the NMR study is consistent with the electron density map (29). In this lipid configuration, the 2’-acyl chain, being partially removed from the bilayer, binds to a hydrophobic cleft of the MA protein. In another recent experimental study, Saad and coworkers characterized a myristoylated MA lattice structure that has some features similar to those in the immature HIV-1 particles and verified protein-protein binding in the MA assembly structure using X-ray crystallography and NMR spectroscopy (38). They also interrogated an alternate PIP2 binding site in that MA lattice using NMR titration with inositol 1,4,5-triphosphate (IP3), the headgroup of the PIP2 lipid.

Unlike a system of integral membrane proteins, the determination of the specific protein-lipid binding properties of a peripheral membrane protein (PMP) like HIV-1 MA remains an outstanding challenge both for experimental and computational studies because of their transient nature (29,39–45). Long-timescale MD simulations are needed to sample PMP binding with a membrane, which are computationally expensive even for a small monomeric PMP (32,46–48). In a previous all-atom molecular dynamics (AAMD) study, using microsecond-long trajectories, the membrane binding mechanism of MA monomers was characterized, including Myr insertion events and the initial membrane response to MA binding (32). An enrichment of PIP2 at the MA-binding site was reported, as it has been suggested several times before (28,49,50). The analyses by Monje-Galvan *et al.* revealed that initial membrane targeting is mediated by the HBR domain, whereas Myr insertion plays an important role in sorting of membrane lipids around the protein binding site and maintaining the stable membrane binding of MA. Interestingly, the cryo-ET density maps suggest that all the Myr groups predominantly remain in the exposed state, while being inserted into the membrane. Therefore, characterization of membrane-bound MA assembly structures employing AAMD simulations requires Myr groups of all the MA monomers stably inserted into the membrane, which makes the present study even more challenging.

To gain molecular-level insights into the altered MA-MA and MA-lipid interactions upon HIV-1 maturation, we applied long-timescale AAMD simulations using cryo-ET fitted atomic model structures of MA assemblies and a realistic asymmetric membrane model. We also performed biased free energy sampling simulations using a combination of steered MD (SMD) and restrained MD (rMD) simulations to sample the conformational space of the MA-bound lipids. Some of the key findings of our study are as follows:

1. Our AAMD simulations verified the stability of MA-MA interactions reported by cryoET data and identified additional interactions in the trimeric interfaces.
2. Simulations explored PIP2/PS binding sites in MA protein complexes. Here we report an alternate PIP2 binding site in the immature MA trimer-trimer interface.
3. Enhanced sampling simulations explore novel conformations of PIP2/PS lipids at the trimeric interface of both complexes and obtain a stable binding of two PIP2 lipid molecules, at extended conformations where the 2’ acyl tail of PIP2 interacts with helix 4 residues of MA, unlike in the immature MA complex.
4. Time-averaged lipid density maps and quantitative analyses of lipid count/fraction data in the vicinity of the MA trimeric interfaces suggest an enrichment of inner-leaflet anionic phospholipids (PIP2) as well as outer-leaflet cholesterol lipids near the MA trimer-trimer interface in the mature MA complex compared to immature MA complex.

The following sections report detailed analyses of our simulated data as well as our simulation methodology.

## Results

### MA trimer-trimer interactions (TTIs) in immature and mature virion states

As discussed earlier, MA trimers in the immature virion get rearranged upon HIV-1 maturation. Our simulations explored atomic-level stable and transient protein-protein interactions between the membrane-bound MA trimers in the immature and the mature MA complexes (**Figure 1** and **Figure 2**). MA monomers consist of five α-helices and one shorter 3_10_ helix between helix2 and helix3 (**Figure S1** in *Supporting Information*). A flexible loop domain, connecting helix1 and helix2, is enriched with basic residues, denoted as HBR. As mentioned earlier, the HBR domain and Myr group play a crucial role in the membrane targeting and stable membrane binding of MA. Myr-MA protein primarily oligomerizes as trimers and further multimerizes in vitro and in virus to form a lattice of hexamer-of-trimers (19,28). To assess trimer-trimer interactions (TTIs) of MA assembly in the immature and the mature virions, we performed AAMD simulations with a dimer of MA trimers (derived from cryo-ET density map (19)) bound to an asymmetric membrane. The study of protein-protein interactions (PPI) in such a peripheral membrane protein complex is challenging mainly because of two reasons: i) we simulate only a part of the continuous lattice structure which renders instability or distortion to the simulated structures, ii) myristoyl insertion of the MA monomers requires long time-scale simulations. However, in both the assemblies, a comparison of the simulated structures and the cryo-ET density map verifies that Myr groups in both immature and mature MA lattice structures remained inserted into the membrane. Although the MA trimer-trimer interface structure is stable during simulations, simulated trajectories captured some deviations of the MA structures from cryo-ET density maps, especially for the membrane-bound N-terminal helix1 in both immature and mature MA models.

**Figure 1:**
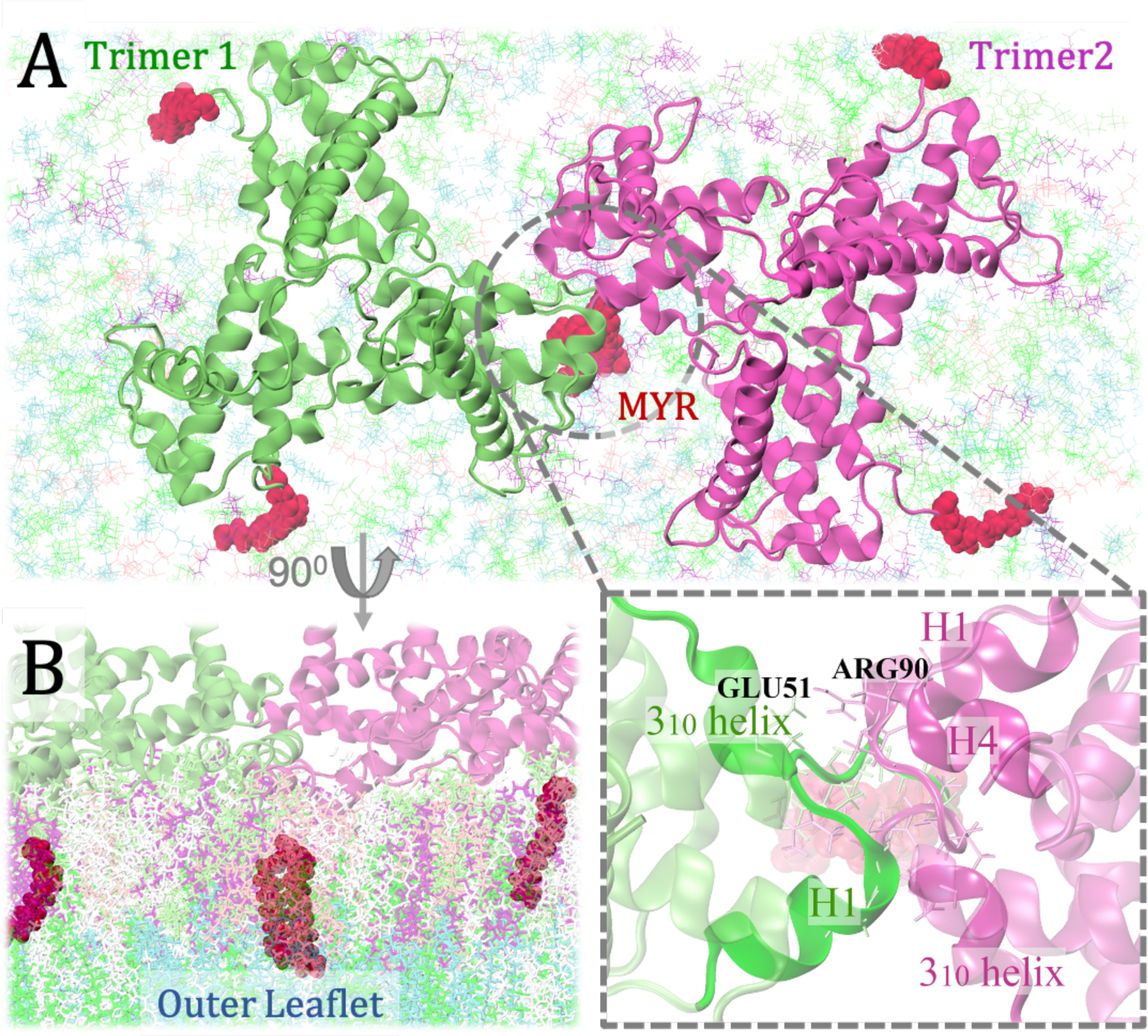
MA trimer-trimer interactions (TTIs) in the immature HIV-1 particles. A, B. Bottom (as viewed from the virus center) and side-views of MA protein assembly (dimer of trimers) with asymmetric membrane. N-terminal Myr groups are inserted into the inner leaflet of the bilayer which is consistent with the cryo-ET density map [EMD ID: 13087]. C. MA domains participating in TTIs are mainly NTD, H1, H4, and 3_10_ helices.

**Figure 2:**
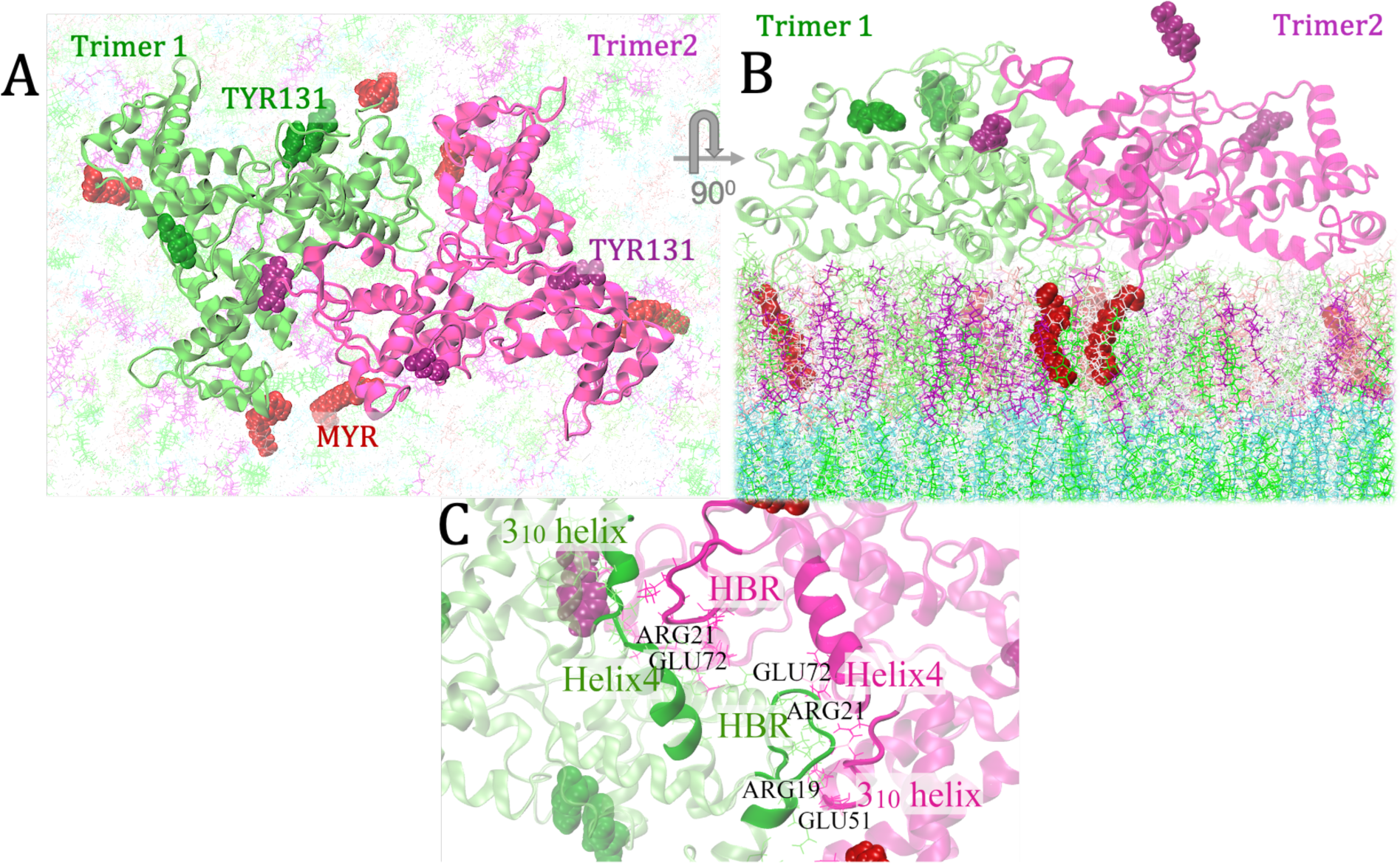
MA trimer-trimer interactions (TTIs) in the mature HIV-1 particles. A, B. Bottom(as viewed from virus center) and side-views of MA protein assembly(dimer of trimers) with asymmetric membrane. N-terminal Myr groups are inserted into the cytoplasmic leaflet of the membrane. C. TTIs are mainly mediated by highly basic region (HBR), H4, and 3_10_ helices. Important residues stabilizing the trimer-trimer interface through electrostatic interactions are highlighted.

Unlike in the mature MA assembly, only two MA monomers, belonging to different trimers, form the trimer-trimer interface in the immature MA assembly. The cryo-ET derived structural model reported that TTIs in the immature MA assembly are mainly mediated by the N-terminal residues and the residues from H1 and 3_10_ helices (19). Throughout our simulations, these interactions remained stable. N-terminal residues of each MA monomer interact with N-terminal residues and H1 helix residues of the other MA monomer in the immature MA assembly. These interactions are mainly mediated by hydrophobic residues ALA2, ALA4, VAL6, and GLY9. Some of the stable contacts explored by our simulations are ALA2:ALA2, ALA2:ALA4, GLY9:VAL6, SER5:GLY9, and ALA4:GLY9. Myr residues of both the MA monomers remain inserted into the membrane in a stable configuration and interact with each other, rendering stabilization to the trimer-trimer interface (***Figure 1*** and **Figure S2**). The N-terminal residues, including ARG3, are observed to offer a stable binding site for PIP2 lipid in the immature assembly which will be discussed later, in detail. Interestingly, in the immature MA assembly, N-terminal residues which assist in the conformational switch during Myr insertion, mediate TTIs. As observed in our simulated trajectories, LEU30 of the flexible HBR loop transiently interacts with hydrophobic residues of NTD of the same and the other MA monomer. Apart from these, our simulations explored a stable electrostatic interaction between ARG90 (the C-terminal residue of H4) of one MA monomer with the acidic residue of 3_10_ helix, GLU51 of the other MA monomer across the trimeric interface. As mentioned earlier, structures of MA monomers at the trimer-trimer interface throughout the simulated trajectories show some deviations from the cryo-ET density map. Interestingly, the membrane-bound helices of MA trimer1 in the immature MA complex (with a bound PIP2 molecule at the alternate binding site) fit better to the cryo-ET density map than the other MA trimer. MA-PIP2 binding is discussed in the next section, in detail.

By contrast, in the mature MA assembly, the reorganization of trimers enables positively charged residues from HBR domains to participate in the trimer-trimer interactions (19). Unlike the immature MA assembly, two monomers of each trimer mediate trimer-trimer interactions. HBR residues of one MA monomer of trimer1 bind to residues of helix4 and 3_10_ helices of the other two monomers of trimer2 (19). In our simulations, arginine residues from the HBR domain (ARG19, ARG21) bind to acidic glutamate residues (GLU51 and GLU72) of 3_10_ and helix4 (***Figure 2*** and **Figure S3**). Unlike the ARG21:GLU72 interaction, our simulations didn’t sample ARG19:GLU51 interaction in both pairs of monomers at the trimer-trimer interface. Our simulations explored additional TTIs mediated by C-terminal cleaved end and Helix 5 residues. In our simulated systems of mature MA, TYR131 of trimer2 (shown as purple beads in ***Figure 2***) binds to ARG57 of trimer1 through stable cation-pi interactions. Apart from ARG57, the binding pocket contains LYS112, GLY61, ILE103, TYR78, ALA114, LEU60, GLN58, GLN115, GLN64. It should be noted that we report here stable MA-MA interactions as sampled by our simulated model systems. However, the protein crowding effect in a mature virion can impact this kind of protein-protein interactions mediated by TYR131 and therefore, this mode of MA-MA interactions might not play an important role in the virion.

Overall, AAMD simulations with the membrane-bound dimer of MA trimers suggest Myr groups of two MA monomers at the dimeric interface, while inserted into the bilayer, interact with each other, thus stabilizing the MA trimer-trimer association in the immature virion. As reported by cryoET data, N-terminal residues and the residues of H1 and 3_10_ helices mediate TTIs in the immature MA assembly. Besides these interactions, primarily hydrophobic in nature, our simulations suggest that electrostatic interactions between GLU51 (3_10_ helix residue) and ARG90 (H4 helix residue) play an important role in stabilizing the trimer-trimer interface of immature MA assembly (***Figure 1***). On the other hand, in the mature virion, two MA monomers from each trimer comprise the inter-trimer interface, and the interactions are mediated mainly by the basic residues of HBR domains and the acidic residues of 3_10_ and H4 helices. The interactions between ARG21 (HBR residue) and GLU72 (H4 helix residue) in both pairs of MA monomers at the trimer-trimer interface were maintained throughout our simulated trajectories (**Figure S3**). Apart from these, GLU51 was observed to interact with ARG19 of the HBR domain in our simulations. The reorganization of MA trimers is presumed to impact the MA-lipid interactions, which will be discussed in the following sections.

### Specific interactions of the protein complexes with PIP2 lipids

Phosphatidylinositol 4,5-bisphosphate [PI(4,5)P_2_ or PIP2], a minor phospholipid component of the plasma membrane (PM), plays a crucial role in the HIV-1 assembly process (24,51–53). A common feature of all eukaryotic plasma membranes (PM) is the asymmetric lipid distribution in its two leaflets, PIP2 being primarily concentrated on the cytoplasmic (inner) leaflet of PM. This lipid molecule contains a negatively charged multivalent inositol head group that remains a few angstroms above the membrane surface. Furthermore, with one saturated tail (C18:0, 1’ acyl chain) and one polyunsaturated tail (C20:4, 2’ acyl chain), this lipid molecule can cause lipid packing defects in the membrane (54).

Long-timescale unbiased AAMD simulations of Myr-MA assemblies bound to the asymmetric model membrane revealed specific lipid binding sites at the interface of two MA trimers, where MA-membrane binding is mediated by the electrostatic interactions of the lipid headgroups with basic amino acid residues. Time-averaged number density profiles of PIP2 headgroup atoms are computed with the last 200 ns of trajectories. The lipid density map reports a low PIP2 lipid density at the trimer-trimer interface of immature MA assembly. An alternate binding site of PIP2 was observed (denoted as ‘a’ in ***Figure 3***) in which a multivalent inositol headgroup interacts with ARG3 of both the monomers at the interface and LYS29, LYS31 of T1:M3 (monomer3 of trimer1) (**Figure S4**). Simulated data agrees with an NMR titration experiment with inositol 1,4,5-triphosphate (IP3) (the headgroup of PIP2 lipid), which reported an alternate group of signals corresponding to N-terminal residues and residues 27-32 for immature-MA like crystal structure (38). The 2’ acyl chain of this PIP2 lipid has been observed to interact transiently with Myr groups at the trimer-trimer interface.

**Figure 3:**
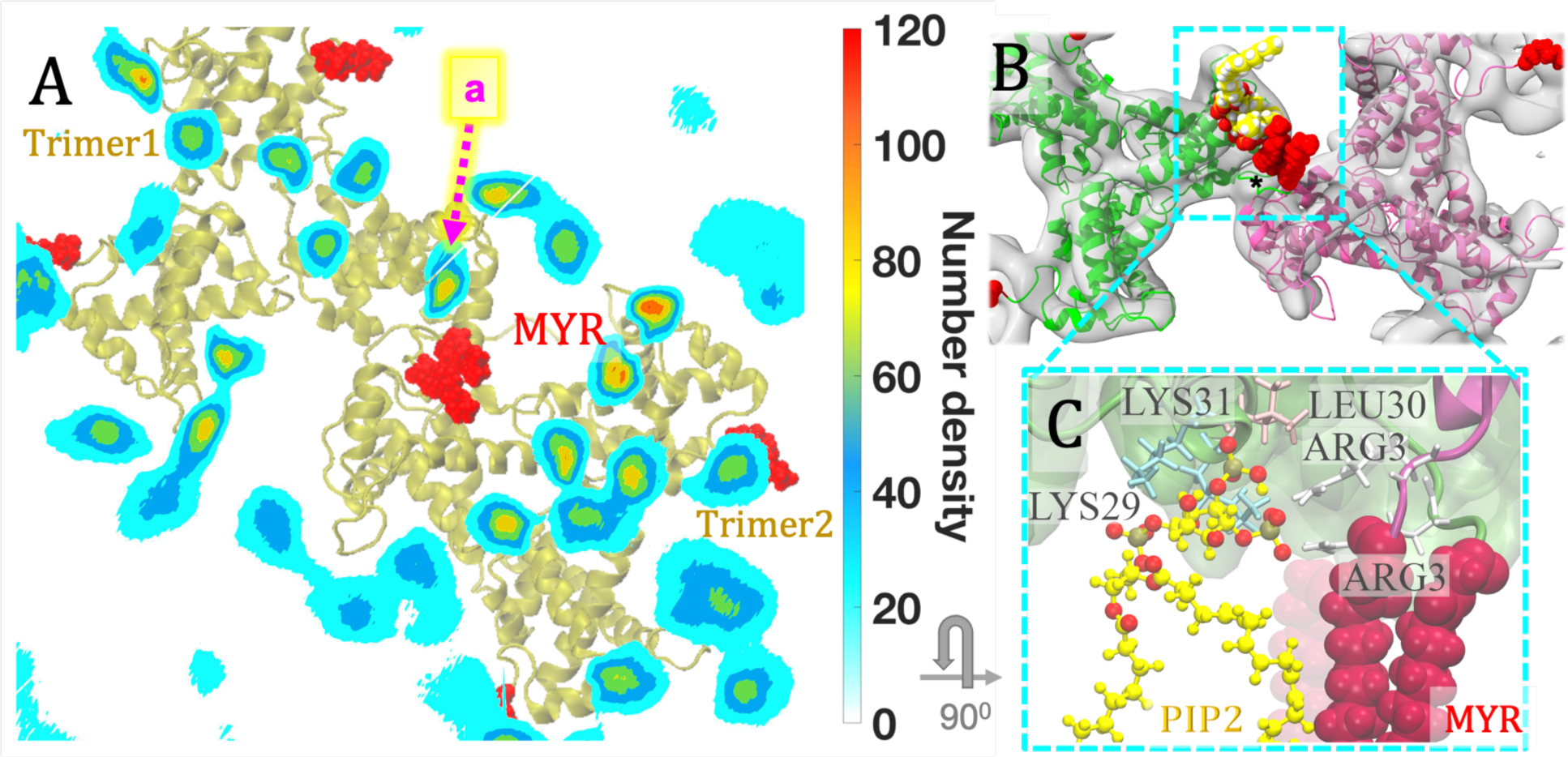
Alternate PIP2 binding site at the immature MA trimer-trimer interface. A. Density map of the PIP2 lipid headgroups in the bilayer plane overlapped with immature MA complex, as viewed from the top towards the virus center. The alternate stable binding site at the trimer-trimer interface is denoted as “a”. B. PIP2 lipid molecule (yellow) at this binding site transiently interacts with Myr groups (red) with its 2’-acyl tail. Cryo-ET density marked with an asterisk corresponds to N-terminal residues linked to Myr groups. C. A close-up view of the residues involved in the alternate PIP2 binding site which includes positively charged LYS29, LYS31 residues of the HBR domain of MA, and N-terminal ARG3 residues of both the MA monomers at trimer-trimer interface.

For the mature MA assembly, our AAMD simulations revealed that the trimer-trimer interface is enriched with PIP2 lipids. Inositol headgroups of PIP2 lipids bind to the charged residues of the HBR domain at the trimer-trimer interface. As discussed earlier, using an NMR approach, Saad *et al.* demonstrated one possible mode of direct binding of MA with a truncated, soluble PIP2 derivative [tr-PI(4,5)P_2_], in which tr-PI(4,5)P_2_ adopts an “extended conformation” with 2’-acyl chain bound to a hydrophobic cleft of MA containing ARG76, SER77, THR81, TRP36 (29). This NMR structure is consistent with the cryo-ET density map [EMD-13088] which contains density components at those PIP2 binding pockets of mature MA (19). Taken together, NMR and the cryo-ET experimental data lead to an important question: Besides the role of PIP2 in membrane targeting and stable membrane binding of MA, does it play a crucial role in HIV-1 membrane maturation?

To examine this hypothesis, we aimed to explore the conformational space of MA-bound anionic phospholipid molecules, mainly multivalent PIP2 and monovalent PS (phosphatidylserine) lipids at the trimer-trimer interface of both the immature and mature MA complexes. To investigate possible “extended lipid conformations”, we performed non-equilibrium steered MD (SMD) simulations with multiple initial structures selected from the last 500ns of 4 μs trajectory for both the complexes [see *Supporting Information* for details]. The center-of-mass (COM) of the selected lipid molecules was pulled in a direction away from the bilayer center with SMD simulations, followed by restrained MD and unbiased MD simulations to examine the lipid conformational stability for these two MA complexes.

Among several selected PIP2 and PS binding sites, PIP2 lipids in two of them (the combined density is denoted as ‘a’ in **Figure 4A**) sampled a stable “extended conformation” in which 2’-fatty acid chain binds to the MA protein while the I’ lipid tail is still inserted into the membrane. In this mode of MA-PIP2 binding, the inositol headgroup of PIP2 interacts with positively charged residues like LYS25, LYS26, ARG21, and 2’ acyl chains of PIP2 molecules, being partially removed from the membrane, binds to helix4 residues such as GLU72, ARG75, SER76, ASN79 (**Figure 4D**). Note that ARG21 residues at the interface are predominantly occupied by interaction with the PIP2 lipid headgroup; however, PS lipids bind to LYS26, LYS29, and LYS31 of the HBR domain (***Figure 8***). None of the PS lipids at the trimeric interface of the mature MA complex sampled such extended lipid conformation. The interactions between MA residues, SER76 and ASN79, interacting with the 2’ acyl tail of PIP2 lipid observed in our simulations agree with NMR predictions (**Figure S5**)(29). Nevertheless, in the simulated trajectories, these PIP2 lipids occupy a different position within the MA trimer-trimer interface from that observed by NMR and cryo-ET. Here we report a pair of PIP2 lipids that bind the HBR domain with their multivalent headgroups, which can access the 2’ acyl tail binding pocket in a stable configuration. However, further simulations and higher resolution cryo-ET data are required to confirm which PIP2 lipid (in terms of head group binding) accesses the 2’ acyl tail binding pocket in the actual virion environment. Similar to the mature MA complex, we also explored the possible extended conformations of the PIP2 lipids close to the trimeric interface in the immature MA complex, which will be discussed in the next section.

**Figure 4:**
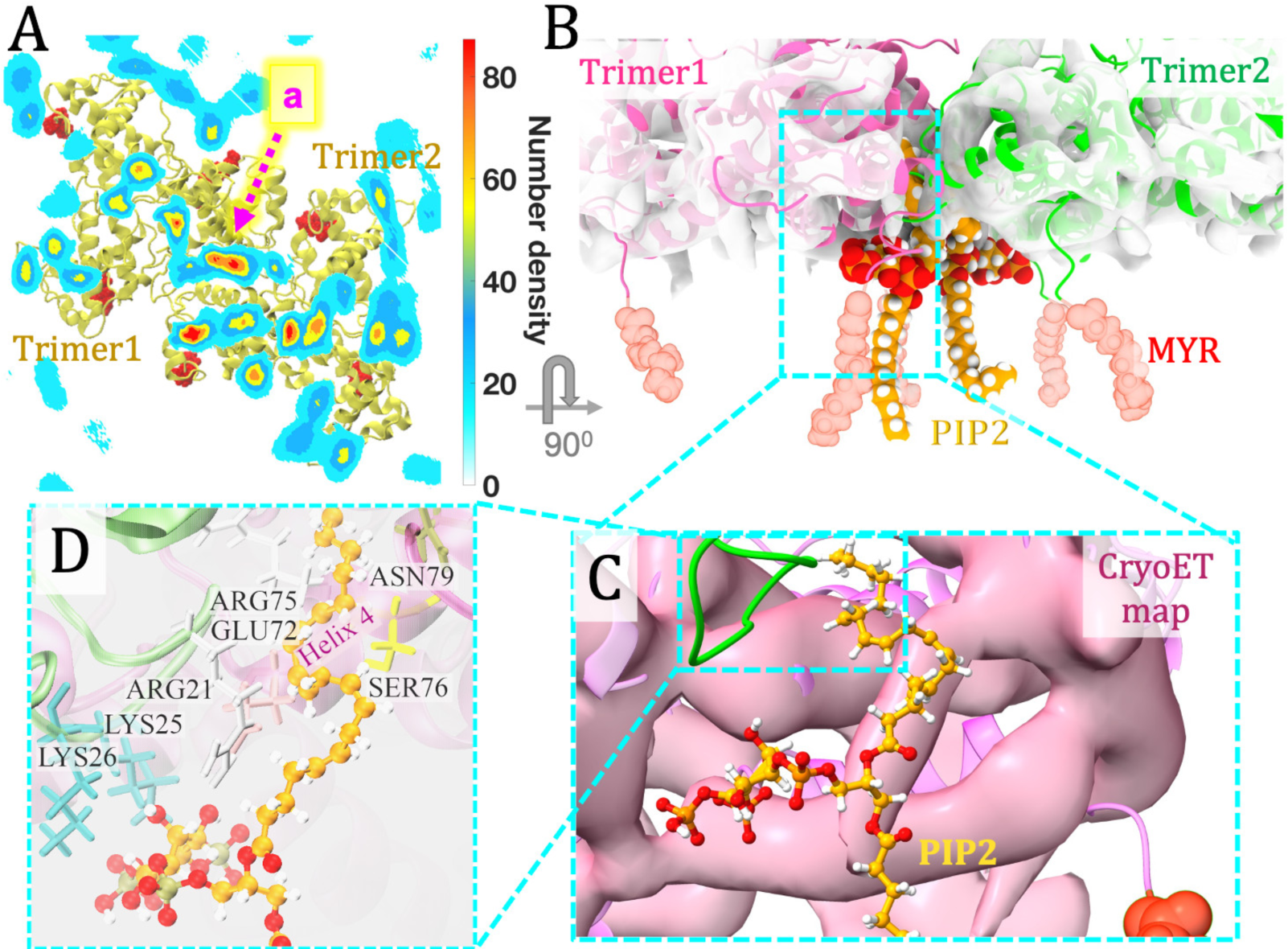
PIP2-binding sites at the mature MA trimer-trimer interface. A. The density map of PIP2 lipids in the bilayer plane. Stable binding sites at the trimer-trimer interface are denoted as “a”, where two PIP2 lipid molecules bind MA proteins with their inositol headgroup, as well as the 2’ acyl chain. B. A close-up view of the PIP2 binding site in the mature MA complex. PIP2 lipid tails and headgroups are shown in orange and red, respectively. C. Simulated MA-PIP2 structure is fitted to the cryo-EM density map of the mature MA complex (Pink; EMD-13088). Although the PIP2 lipid, especially its 2’ acyl tail, is close to the cryo-ET observed density, they do not superimpose with each other. D. The residues at the mature MA-PIP2 binding site, conferring stability and specificity to the complex are ARG21, LYS25, LYS26 (bind the headgroup of PIP2), GLU72, ARG75, SER76, ASN79 (bind the 2’ acyl chain).

### Simulated MA-PIP2 interactions are consistent with previous photocrosslinking data

In a previous set of experiments, a functionalized PI(4,5)P_2_ derivative (f-PI(4,5)P_2_) was incorporated in the immature and mature virus particles to study MA-PIP2 interactions (19). The f-PI(4,5)P_2_ contains a diazirine group covalently linked to its 1’ acyl chain that can be photo-crosslinked to nearby MA protein. Upon photoactivation, f-PI(4,5)P2 was observed to be efficiently crosslinked to MA within immature particles, but not in mature particles.

Our AAMD simulations explored extended conformations of PIP2 lipids in the immature MA complex and probable binding pockets of MA. Our findings reveal that, in immature MA assembly, unlike mature MA, the stable extended conformation of PIP2 has its 1’ acyl chain binds to MA while partially removed from the membrane. Simulated trajectories sampled two binding pockets, one of them (Binding pocket 2 or BP2 as shown in **Figure 5**) contains acidic glutamate residues(GLU72, GLU73) which are able to be crosslinked to f-PI(4,5)P_2_ upon photoactivation(55). The other binding pocket is enriched with hydrophobic residues, such as VAL83, TYR28, TRP15, LEU30, LEU20, and THR80. For both the binding modes, the inositol headgroup of the PIP2 lipid binds to ARG3 of both the MA monomers at the trimeric interface and LYS29. **Figure 5G** shows the switch of PIP2 lipid between two binding pockets as obtained from an unbiased MD trajectory. 1’ acyl chain binding to the hydrophobic cleft (VAL83:BP1) is observed to be more probable than binding to binding pocket 2. Further, in the mature MA structure, GLU72 and GLU73 are occluded by the 2’ acyl tail of bound PIP2 (**Figure 4**). This provides a possible explanation for the experimental observation that f-PI(4,5)P2 was efficiently crosslinked to MA upon photoactivation within immature particles, but not to MA within mature particles.

**Figure 5:**
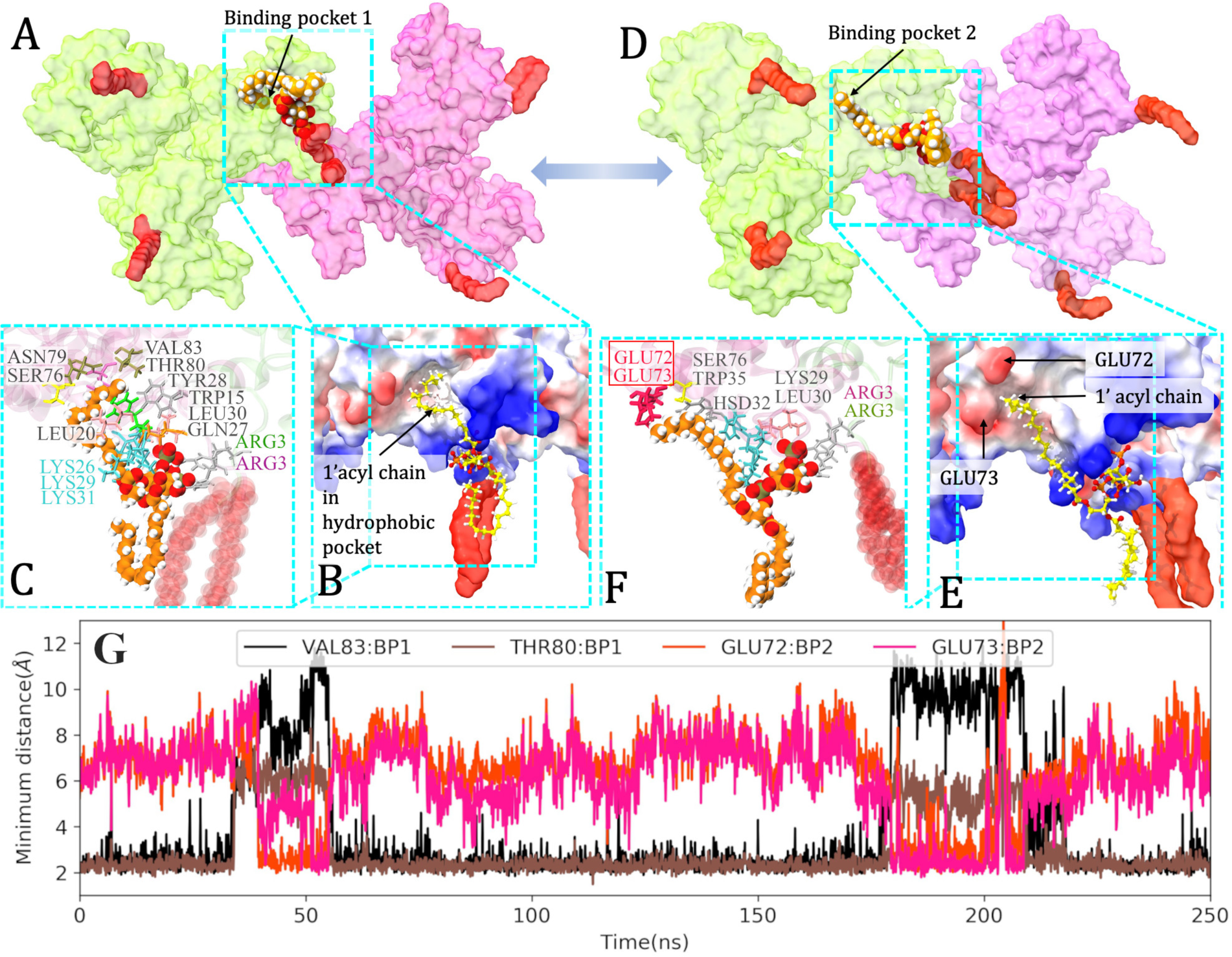
Accessible binding pockets for 1’ acyl chain of PIP2 in immature MA complex. (A-C). Binding pocket 1(BP1) contains hydrophobic residues, VAL83, TYR28, TRP15, LEU30, LEU20. (D-F) Binding pocket 2(BP2) contains acidic GLU residues (GLU72, GLU73), shown in red in (F), which preferentially reacts with alkyl diazirine moiety attached to 1’ acyl chain of f-PI(4,5)P2 (55). B, E. Electrostatic surface potential map of immature-MA protein. Red and blue colors represent negatively and positively charged electrostatic potential, respectively. G. Minimum distance of 1’ acyl chain of the PIP2 lipid in an extended conformation, from VAL83, THR80 residues in BP1 and GLU72, GLU72 residues in BP2.

In summary for this section, our enhanced sampling simulations explored stable binding pockets for PIP2 lipid tails close to the immature and mature MA trimer-trimer interface. In the mature MA assembly, the 2’ acyl tail of PIP2 lipid is observed to sample a stable binding pocket of MA consisting of helix4 residues such as GLU72, ARG75, SER76, and ASN79. In contrast, in the immature MA assembly, the 1’ acyl tail sampled two binding pockets while the headgroup of the PIP2 lipid binds to an alternate site containing N-terminal residues and HBR residues such as LYS29 and LYS31. These results provide a possible structural explanation of how PIP2 lipids bind immature and mature virions with different binding modes.

### Lipid sorting around MA complexes

Next, we investigated the sorting of other membrane lipids around the HIV-1 MA trimer-trimer interface in the immature and mature virus particles. Although HIV-1 immature particles bud from the plasma membrane (PM) of the infected cell, the lipid composition of the viral particle membrane differs significantly from the host cell PM. However, like the PM, the HIV-1 membrane is also asymmetric in lipid distribution and lipid unsaturation. The inner leaflet is enriched in unsaturated phospholipids, like phosphatidylethanolamine (PE), phosphatidylserine (PS), phosphatidylcholine (PC), phosphatidylinositol (PI) lipids, and cholesterol, whereas the outer leaflet mainly contains phosphatidylcholine (PC), saturated sphingomyelin lipids (SM), and cholesterol. Experimental studies suggest that during the HIV-1 assembly process, Gag multimerization occurs in the specialized microdomains of the inner leaflet of PM, enriched in PIP2 and cholesterol (21,22,25,56). Sphingomyelin and cholesterol in the outer bilayer form densely packed liquid-ordered phases, i.e., “lipid rafts”, while PIP2 in the inner bilayer prefers the liquid-disordered phase due to its unsaturated tail.

***Figure 6*** presents the time-averaged lipid density maps for PS in the inner leaflet, cholesterol in both the leaflets, and SM lipids in the outer leaflet for both the MA complexes. Number density profiles of lipid headgroup atoms are computed with the last 200 ns of the trajectories. As already discussed, MA trimer reorganization is observed to alter specific interaction sites of PIP2 lipids (***Figure 3*** and ***Figure 4***). Also, for PS lipids, the computed lipid density profiles show high-density domains at the trimer-trimer interface of the mature MA complex, especially close to the HBR loops. The headgroups of acidic PS lipid molecules bind to the basic lysine residues of the HBR domains, whereas specific binding sites of PIP2 lipid molecules include ARG21 of the two HBR domains (**Figure 8**). On the other hand, the trimer-trimer interface of the immature MA complex contains a lower number of high-density domains of PS lipids compared to the mature MA assembly (***Figure 6A,B***). In the computed lipid density profile for the immature MA assembly, PS lipid density is observed near the HBR domain, however, those interactions are not maintained throughout the trajectories as captured by contact frequency calculation (**Figure 8**). Also, PS lipid molecules are observed to interact with N-terminal residues in the immature MA assembly. On the other hand, PIP2 lipid molecules bind the HBR domains of trimer1, trimer2, and an alternate binding site consisting of N-terminal residues of both the trimers and the HBR domain of trimer1 (***Figure 3*** and **Figure 8**). We further looked into the distribution of PS lipids around PIP2 lipids in the mature MA assembly and studied the effect of MA-membrane binding on such distribution. The radial distribution functions shown in **Figure S6** suggest an enrichment of PS lipids around PIP2 lipids at the specific binding sites of the MA trimer-trimer interface of mature MA complex, compared to the other PIP2 lipids in the simulated system.

**Figure 6:**
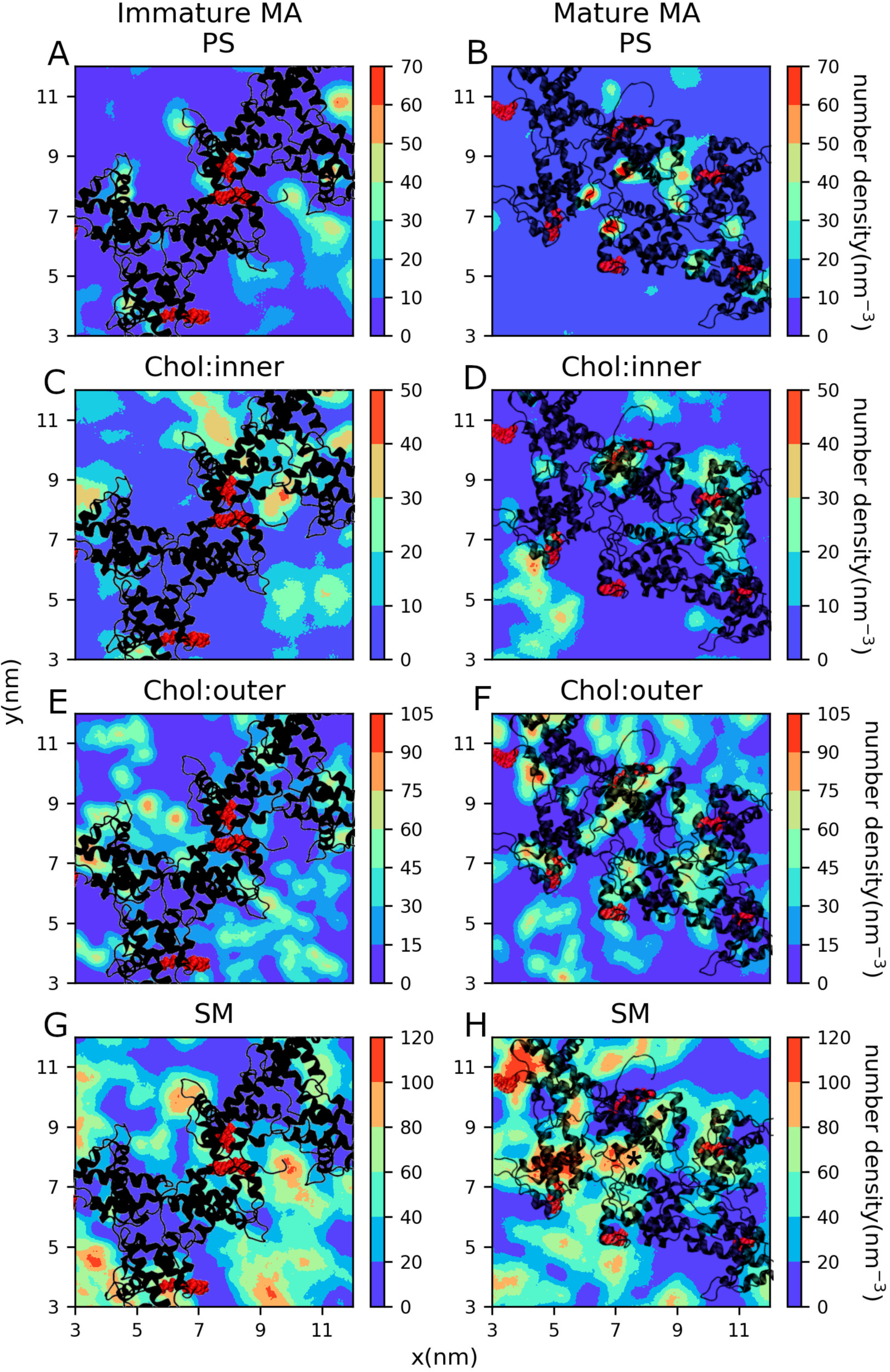
2D-density maps of lipid bilayer species at the trimer-trimer interface of the immature and mature MA assembly. (A-B) Phosphatidylserine (PS) in the inner leaflet, (C-F) cholesterol in the inner and outer leaflets, and (G-H) sphingomyelin(SM) in the outer leaflet. MA complex (as viewed from the virus center) is shown in black with Myr groups highlighted in red.

Cholesterol molecules constitute a major population (up to ∼30%) of the total species in the HIV-1 viral membrane (21,22), which helps counteract unfavorable packing of saturated and unsaturated lipids in membranes in general and maintain structural integrity and membrane fluidity. Time-averaged density maps for raft-forming cholesterol and SM in the outer leaflet, obtained from our simulated AAMD trajectory, exhibit high-density domains of those components near the MA complexes. In the past, multiple studies proposed that outer leaflet rafts could be trapped by inner leaflet PIP2/PS-Gag nanodomains through the trans-bilayer coupling (34,57–59). Our trajectories show transient interactions between inner-leaflet MA-bound anionic phospholipids and outer-leaflet SM, however, the contribution of inter-leaflet coupling and protein-mediated lipid raft formation in such systems is beyond the scope of the present study and needs to be carefully examined in the future by computational and experimental studies.

### Quantitative characterization of MA-lipid interactions and lateral organization of the membrane

We examined the MA-lipid interactions and lipid arrangement in the vicinity of MA complexes, in further detail. In the last section, time-averaged lipid density maps are reported. The high-density domains in those maps may occur either by a greater number of lipid molecules in that particular domain or the dynamic localization of lipids. Here we have computed the number/fraction of lipid molecules around MA monomers at the trimeric interface (TI) of both immature and mature MA complexes to compare their lipid environment quantitatively, as captured in our simulations. Here we have used the last 200 ns of the trajectories, similar to the lipid density profile calculation (**Figure 6**). For the PIP2 and PS lipid count calculation, we have considered a distance cutoff of 10 Å between the phosphorus (P) atoms of phospholipid headgroups and the Cα atoms of the MA monomers adjacent to the TI. **Figure 7A** and **Figure 7C** show the probability of PIP2 and PS lipid count values for immature and mature MA trimer-trimer interfaces, respectively. According to the histogram data, the maximum probability is observed for 2-3 PIP2 and 3 PS lipids close to the two MA monomers in the immature MA TI and 8-9 PIP2 and 5 PS lipid molecules close to the four MA monomers in the mature MA TI. Therefore, the number of MA-PS interactions per MA monomer at TI is not different in these two MA complexes, as sampled in our simulations. However, these data suggest the binding of more PIP2 lipids to the MA monomers at the TI in the mature MA complex. The fluctuation in the lipid count data depends on the measurement criteria and the dynamic MA-lipid interactions. However, at a time instant, the number of PIP2/PS lipids in the vicinity of MA monomers at the TI should follow these statistics. We further investigated the binding frequency at different interaction sites to determine the stability of those interactions, which will be discussed later in this section.

**Figure 7:**
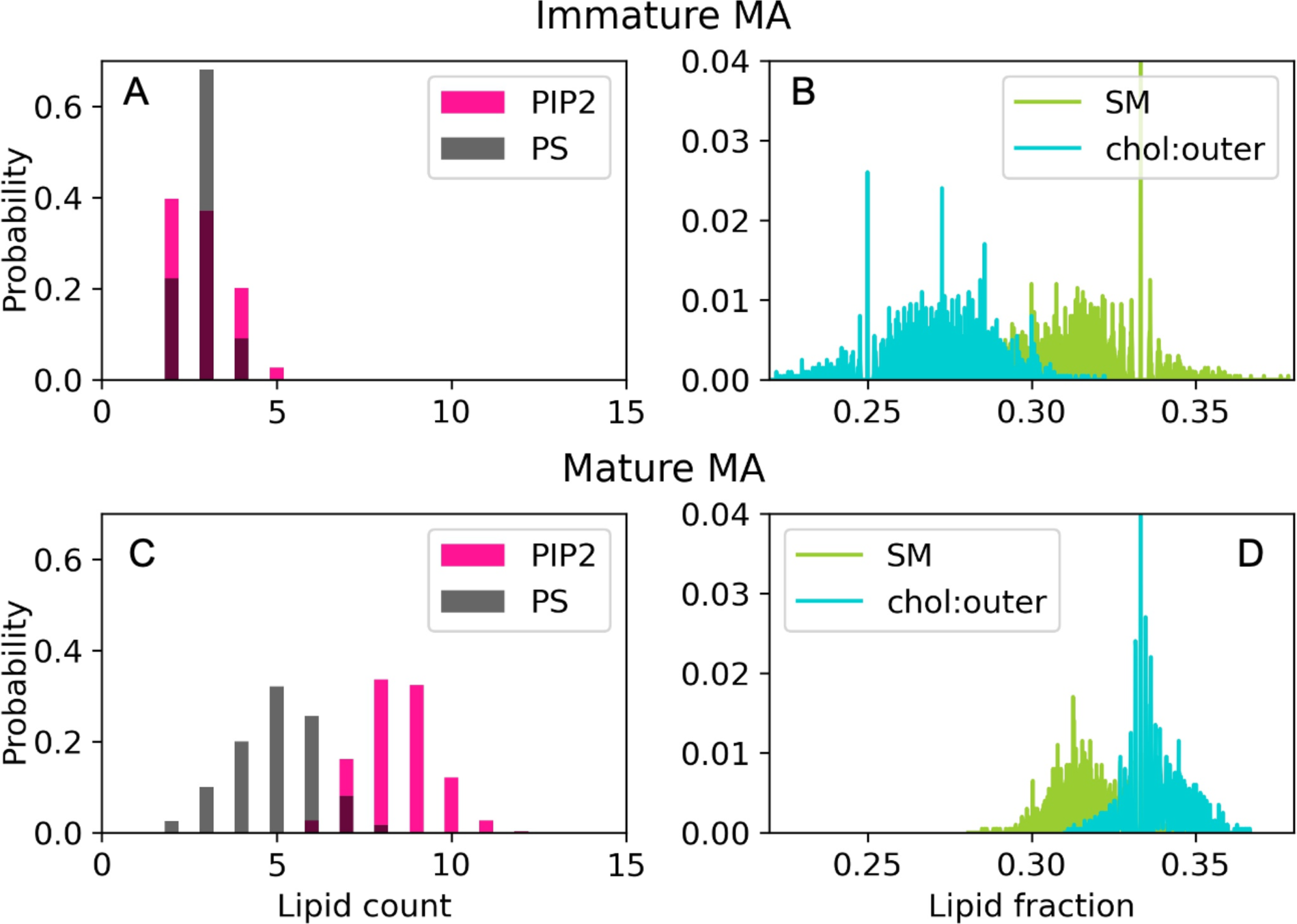
Probability distribution of the number of lipid molecules/fraction of lipid molecules below the trimer-trimer interface of (A-B) the immature MA and (C-D) mature MA complexes.

For the calculations of SM and cholesterol lipid count at the outer leaflet of the bilayer, we have considered different parameters and selection criteria. As the number of MA monomers at the TI is different, the surface area under those MA monomers is considerably different. Therefore, instead of lipid count, here we compare the fraction of lipid molecules near the MA monomers with respect to the total number of lipid molecules in the selected domains. Using the same trajectory segments, the number of SM/Chol lipids at the outer leaflet is computed using a distance cutoff of (1) 55 Å between the Cα atoms of MA monomers and the P atoms of SM lipids, and (2) 50 Å between the Cα atoms of MA monomers and oxygen atoms of the hydroxyl group of outer-leaflet cholesterol molecules. **Figure 7B** and **Figure 7D** present the probability distribution of the fraction of SM/cholesterol lipid near the MA monomers at the TI of immature and mature MA complexes, respectively. Although the fluctuation of the fraction of SM lipids is more in the immature MA complex, the probability distribution profiles of SM lipids in the immature and mature MA complexes overlap with each other. However, we observe significantly different distribution profiles for the fraction of outer-leaflet cholesterol. This result suggests an enrichment of cholesterol lipids in the outer leaflet near the MA monomers at the TI of mature MA complex, compared to that of immature MA complex. It should be noted here that a single MA monomer or MA trimer also contributes to the lipid enrichment, however, both of these contributions are present in immature and mature MA complexes and by comparing these two systems one can determine the effect of trimer-trimer interactions on the MA-membrane interactions for the MA monomers at the TI. The trend of lipid binding/sorting data presented in **Figure 7** has been reproduced using multiple simulation trajectories and different time domains of a particular trajectory. In our membrane model, long-chain C24:0 sphingolipids in the outer leaflet can interact with inner-leaflet PS/PIP2 lipids. Stable MA-PIP2/PS interactions can cause localization of such lipids compared to other PS/PIP2 lipids in the system with higher lateral mobility. This can impact the dynamic organization of the outer-leaflet SM lipids and create a high-density domain of SM lipids beneath the mature MA trimer-trimer interface as observed in **Figure 6H**. However, quantification of such interdigitation is beyond the scope of this current study.

Next, we examined the strength of PIP2/PS binding at different sites of MA monomers at the TI of immature and mature MA complexes by computing binding frequency. For this purpose, we have used a distance cutoff of 2.5 Å for the minimum distance between MA residues and PIP2/PS lipids. The data presented in ***Figure 8(A,E)*** suggest that N-terminal basic residues (ARG3), HBR residues (ARG19, ARG21, LYS25, LYS26, LYS29, LYS31), and helix2 residues (ARG38, ARG42) exhibit stable interactions with PIP2 in both systems. Most of these residues show a contact frequency of >100% for PIP2 binding which signifies more than one PIP2 lipids bind/remain in contact (within the cutoff distance) with a single amino acid residue. Unlike this, MA-PS binding frequencies suggest stable binding of only one PS with basic residues of MA. As we discussed earlier, our simulations revealed an alternative binding site for PIP2 at the TI of immature MA complex, that contains HBR (LYS29, LYS31) residues and ARG3 of monomer1 (M1) and ARG3 of monomer2 (M2) (as shown in ***Figure 3***). The contact frequency data verify the stability of such binding (**Figure 8A**). In the immature MA complex, most of the PS lipids are observed to interact transiently with MA residues (***Figure 8C***) and the contact frequency data for two MA monomers change at different segments of our trajectories. On the other hand, our simulations sampled stable binding between PS lipids and HBR residues in the mature MA complex (***Figure 8G***). ***Figure 8B, D, F, H*** present snapshots of the MA-bound PIP2/PS lipids and the corresponding *Cα* atoms of MA protein involved in those protein-lipid interactions. Please note that our simulations didn’t attain symmetric MA-lipid interactions for all the MA monomers at the trimer-trimer interface in both systems. For such a peripheral membrane protein complex, the attainment of perfect lipid sorting behavior by MA protein complexes demands a much longer simulation timescale. However, the computation of binding frequency allows us to distinguish between stable and transient interactions, explored in our simulations.

**Figure 8:**
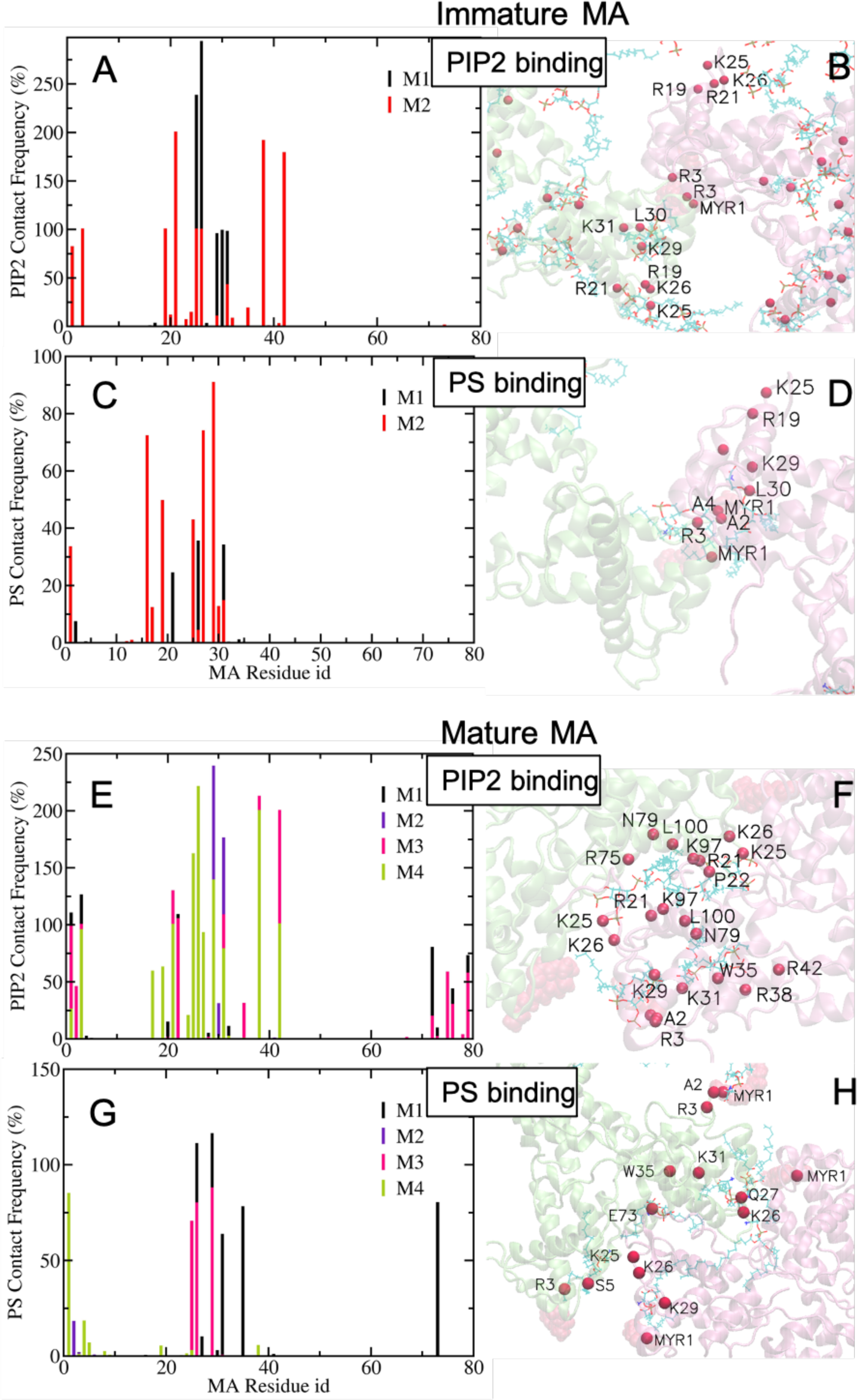
Frequency of PIP2 and PS interactions with each residue of the MA monomers(M1-4) at the trimer-trimer interface of the immature (A, C) and the mature (E, G) MA assembly. Snapshots show the MA-bound PIP2 and PS lipids and MA residues interacting with them in the immature (B, D) and mature (F, H) MA assembly. The Cα atoms of those MA residues are highlighted with red spheres.

Altogether, the long-time-scale MD simulations distinguished the lipid-sorting around the MA trimer-trimer interface in the immature and the mature MA complexes. As captured in the time-averaged lipid density map, the mature MA trimer-trimer interface is observed to be enriched with anionic phospholipids such as PIP2, compared to the immature MA trimer-trimer interface. The enrichment has been quantified by computing the probability of the number of bound-PIP2/PS lipids at the trimeric interface. The contact frequency data analyzed the stable and transient MA-PIP2 and MA-PS interactions. The stability of MA-PIP2 binding at the alternative binding site is verified using the contact frequency data (***Figure 8A*** and ***Figure 3***). Further, the lipid fraction data revealed enrichment of cholesterol at the outer leaflet below the MA monomers at the trimeric interface of the mature MA complex compared to the immature MA complex.

## Discussion

In this work, we have presented a detailed structural analysis of the membrane-bound HIV-1 MA protein complexes as found in the immature and the mature virions. The role of plasma membrane PIP2 lipids in the HIV-1 assembly process is a well-recognized (22,37,52); however, its function in the maturation process is still unknown. Using atomistic molecular dynamics simulations here we interrogated altered MA-lipid-specific interactions and lateral organization of lipids in the vicinity of MA complexes. Different modes of direct binding of PIP2 lipids with HIV-1 MA have been reported by several experimental techniques including NMR (29,38) and mass spectrometric protein footprinting analysis (36). In this study, we have identified the favorable MA-PIP2 binding modes and specific PIP2 binding sites in the immature and mature MA complexes. The binding affinity and arrangement of PIP2, PS, and raft-forming sphingomyelin (SM), cholesterol at the two different trimer-trimer interfaces were quantified using different computational analyses.

The cryo-ET density map captured different organizations of MA trimers in the immature and mature MA lattice structures (19). The electrostatic surface potential of MA proteins facing the central hole of hexamer-of-trimers in the lattice was observed to be altered from positive (in immature MA lattice) to neutral or negative in the mature MA assembly. In addition, the hole becomes smaller in the mature MA lattice. Previously, these holes have been hypothesized to mediate interactions of MA with the C-terminal tail of the HIV-1 Env protein and promote Env incorporation into the virions (27,60), however, the recent cryo-ET experiment suggested that the trimeric Env CTD does not lie in the central hole formed by the immature Gag lattice but instead binds to its hexameric rim, at the MA trimer-trimer interface (20). On the other hand, in the mature virions, the MA layer was observed to be fragmented and Env trimers were observed to gain significant mobility after structural rearrangement.

In the present study, MD simulations with the MA assembly structures derived from cryo-ET density maps explored molecular-level MA-MA interactions at the trimeric interface in immature and mature virions. The characterization of protein-protein interactions remains a challenge for computational studies (61–63). Here we start our simulations from cryo-ET structures of MA multimers and by performing strategic simulations we obtain stable membrane-bound dimer-of-trimer structures of mature and immature MA complexes. We have studied the stability of MA-MA interactions at the trimeric interfaces predicted by cryo-ET structures and explored other possible interactions which might play a role in rendering stability to the protein complexes. As cryo-ET structures reported (19), in the immature MA complex, trimer-trimer interactions are mainly governed by the N-terminal residues, helix1, and 3_10_ helix residues. On the other hand, in the mature MA complex, MA trimer-trimer interactions are mediated by the basic residues of HBR domains and the acidic residues of 3_10_ and helix4. All the crucial MA trimer-trimer interactions reported by cryo-ET data remained stable in our long-timescale simulations. Further, our simulated data revealed that Myr groups at the trimer-trimer interface of immature MA complex, being inserted into the membrane, remain in contact with each other and the hydrophobic interactions between them render stability to the trimeric interface (***Figure 3***). Apart from the hydrophobic interactions mediated by N-terminal residues and H1 residues, our simulated data suggests electrostatic interactions between ARG90 of helix4 and GLU51 of 3_10_ helix stabilize MA trimer-trimer interactions in the immature MA complex (***Figure 1*** and **Figure S2**). A comparison of the structures of MA monomers in our simulated trajectories with the cryo-ET-derived atomic structure shows deviations, especially for the membrane-bound helices of MA monomers at the trimeric interface. Interestingly, the structural deviation for the membrane-bound helices in the MA monomer at the trimeric interface of the immature MA complex is lower when a PIP2 lipid headgroup binds to the alternate site shown in ***Figure 3***. This indirectly verifies this mode of MA-PIP2 interaction in the immature virion. In the mature MA complex, our simulations explored stable electrostatic interactions between pair of residues, ARG19:GLU51. Apart from that, cleaved end residues and more flexible H5 helix residues (LYS97 and LYS113) are observed to participate in the trimer-trimer interactions in our simulated MD trajectories of mature MA complex. Our simulations have identified one stable binding pocket for the post-cleavage C-terminal end residue (TYR131), which contains ARG57, LYS112, TYR78, and other hydrophobic residues such as ILE103, ALA114, and GLY61. TYR131 is able to interact with the basic residue ARG57 through cation-pi interactions. However, in the crowded environment of the mature virion, cleaved end residues may behave differently. Further high-resolution cryo-ET data of mature MA is needed to understand the behavior of cleaved end residues. In summary, the MA trimer-trimer interface changes its nature significantly, as it involves more charged residues upon maturation. This inevitably modifies lipid binding sites across the MA trimeric interface upon MA maturation.

Our long-timescale AAMD simulations characterized the lipid binding to the MA monomers at the trimer-trimer interface of both the MA complexes. Unbiased and biased MD simulations have explored the conformational space of MA-bound anionic phospholipids and obtained stable extended conformations for PIP2 lipid in the mature MA assembly, in which the 2’ acyl chain of the PIP2 molecules is outside the hydrophobic membrane core and is bound to the protein.

Although some of the MA residues (SER76, ASN79) interacting with the 2’ acyl tail of PIP2 lipid in the extended conformation in our simulations match NMR data (***Figure 4*** and **Figure S5**), the overall position of the PIP2 lipid is distinct from the positions as previously suggested by NMR or EM experiments. However, the binding sites of two PIP2 lipids in the extended conformations, as obtained in our simulations, are also at the MA trimer-trimer interface close to the position of the cryo-ET density, as well as the NMR structure. Thus, our simulations suggest that the MA trimer-trimer interface, as in the mature virion, can stabilize extended conformations of PIP2 lipid where the 2’ acyl chain being partially removed from the bilayer can interact with the MA protein. Cryo-ET density map of mature MA lattice predicted the partial removal of up to ∼2500 PIP2 lipids in the mature virion (19). However, the driving force for such PIP2 removal and its impact on the membrane structural and mechanical properties in the mature virion is still an open question. Further, the difference between the positions of PIP2 lipid as obtained in our simulations and cryo-ET data surmise that there may be other stabilizing interactions that are not well represented in the simulations.

Interestingly, this mechanism of MA-PIP2 binding is not operational for the immature MA assembly, but simulations revealed two specific binding pockets of MA for the 1’-acyl chain of PIP2 in the immature MA trimer-trimer interface. One of them is the hydrophobic cleft containing VAL83, THR80, and another contains the acidic residues GLU72, and GLU73 (***Figure 5***). These results provide a possible explanation for the experimental observations using the photocrosslinking approach (19). Functionalized PIP2 derivative [f-PI(4,5)P_2_] with a diazirine group covalently linked to the 1’ acyl chain was cross-linked to the immature MA assembly, but not to the mature MA lattice. A recent study has shown the reactivity of such a diazirine moiety with acidic amino acids such as GLU (55), which are present in one of the binding pockets of MA sampled by 1’ acyl chain of PIP2 in the simulated trajectories of immature MA assembly. On the other hand, these GLU72 and GLU73 residues are preoccupied with the 2’ acyl chain of PIP2 in the mature MA assembly (***Figure 4***). Thus, they cannot bind the 1’ acyl tail of f-PI(4,5)P_2_ in the photo-crosslinking experiment. Therefore, our AAMD simulations add to the observations of photo-crosslinking experiments and suggest an altered binding mode of PIP2 in the mature MA complex.

Further, we quantified the binding affinity of MA-PIP2/PS interactions and verified the stable binding sites for the PIP2 and PS lipids including the alternative binding site of PIP2 at the trimeric interface of immature MA complex (***Figure 8A*** and ***Figure 3***) revealed by our simulations. Our lipid count/fraction data suggest that the simulated system of mature MA trimer-trimer interface is enriched in the anionic phospholipids, PIP2 on the inner leaflet, and cholesterol on the outer leaflet of the bilayer, compared to the MA trimeric interface of immature MA complex(***Figure 7***).

## Conclusion

In summary, we have performed long-time and large-scale AAMD simulations of HIV-1 Myr-MA dimer-of-trimers bound to the inner leaflet of an asymmetric membrane model. We have considered two different Myr-MA structures as reported for immature and mature virions by cryo-ET experiment. A series of carefully performed simulations explored the MA-MA and specific MA-lipid interactions at the interface of MA trimers when Myr groups of all MA proteins remain inserted into the membrane. Our simulations explored electrostatic interactions between ARG90 of helix4 and GLU51 of 3_10_ helix and hydrophobic interactions between Myr groups providing additional stabilization to the immature MA trimeric interface, besides the hydrophobic interactions mediated by N-terminal residues. Here we report an alternative PIP2 binding site in the immature MA complex where the headgroup of multivalent PIP2 lipid binds to ARG3 residues of both MA monomers at the trimer-trimer interface and the LYS29, LYS31 residues of the HBR domain of one of the monomers. We further quantified the stability of such interactions and other MA-PIP2/PS interactions by contact frequency data. Our time-averaged lipid density maps and quantitative calculations of the lipid environment below the MA monomers at the trimeric interfaces of both the complexes suggest an enrichment of PIP2 at the inner leaflet and cholesterol at the outer leaflet of the membrane in the mature MA system, compared to the immature MA system. We explored PIP2 conformational change at the specific binding sites of MA in immature and mature MA complexes using biased MD simulations. Simulated trajectories explored stable binding pockets for the 2’ acyl chain of PIP2 at the trimer-trimer interface of mature MA complex, unlike immature MA complex. In the immature MA complex, our enhanced sampling simulations sampled two binding pockets for 1’ acyl chain, one of them contains hydrophobic residues VAL83, THR80, and another binding pocket contains the acidic residues GLU72, and GLU73. This finding provides a possible structural explanation for the observation of the photocrosslinking experiment. Altogether the present simulation results, combined with prior experimental observations, provide molecular-level insights into the altered binding site and binding mode for PIP2 lipids upon MA maturation. Therefore, specific MA-PIP2 interactions are not merely important for the stable MA-membrane binding during the viral assembly process, it shows interesting characteristics in the virions before and after the maturation process. However, whether the maturation process of MA protein is aided by such PIP2-specific interactions is beyond the scope of our current study. As such, this study reports important insights into altered MA-MA and MA-lipid interactions and fills a significant gap in our understanding of the HIV-1 matrix and membrane maturation.

## Materials and Methods

### All-atom models of the MA trimeric assemblies

The initial all-atom protein configuration was generated by fitting an atomic model of MA trimer (PDB ID: 1HIW) to the cryo-ET density maps, EMD-13087 and EMD-13088, for the immature and mature MA lattice structures, respectively. The monomeric MA structure (PDB ID: 1UPH) (9) with all MA residues up to TYR132 was then superimposed to MA assemblies to obtain the final dimer-of-trimers MA structures to simulate. The Myr group, in an exposed conformation, was covalently attached to its N-terminal GLY1 residue using CHARMM-GUI input generator (64). In the simulated model, residue 1 is composed of the Myr group and GLY residue. Therefore, the residue_n_ of the experimental PDB structure (PDB ID: 1UPH) corresponds to residue_n-1_ of the simulated models. A fully hydrated bilayer was built and equilibrated using CHARMM-GUI Membrane Builder and Quick-Solvator, respectively (64–68)(*see Supporting Information for details*). Next, protein coordinates were positioned close to the inner leaflet of the membrane using VMD software (69) and those lipid molecules, overlapping with Myr groups were removed. The protein-membrane merged structures for both the immature and mature MA were solvated in 150 mM aqueous KCl solution with initial box lengths of 15 nm × 15 nm × 25 nm to allow enough water layers to be present between the protein units and the periodic image of the outer leaflet. The total size of both systems was ∼580,000 atoms.

### Membrane model and protein-membrane system setup

The membrane model designed for this study was based on lipidomic analysis of HIV-1 particles produced in HeLa cells reported by Lorizate *et al*. (32,56). The approximate composition of the extracellular leaflet was 35% cholesterol, 30% phosphatidylcholine(PC), and 35% sphingomyelin(SM) and the cytoplasmic leaflet was 15% cholesterol, 40% PC, 15% PE, 15% phosphatidylserine(PS) and 15% phosphatidylinositol (PIP2). This model mimics the asymmetric nature of the HIV-1 membrane and allows us to examine the effect of trans-bilayer interactions that become operational upon protein binding. The asymmetric bilayer with a surface area of 15 nm x 15 nm was built using CHARMM-GUI Membrane Builder and Quick-Solvator (64–67,70) following a multistep minimization and equilibration protocol. This bilayer system was further equilibrated for 400ns before merging the equilibrated protein coordinates using VMD software (69). All the data shown in the manuscript were computed for the membrane system with DOPC, POPE, DOPS, SAPI, LSM, and cholesterol. However, we have verified MA-MA and MA-lipid interactions with another membrane model with POPC, POPE, POPS, SAPI, LSM, and cholesterol. It should be noted here that the PIP2 concentration in our model membrane has been chosen to be higher than the HIV-1 membrane deliberately so that it allows us to study specific MA-PIP2 interactions within the simulation timescale. Despite that, 4 µs trajectories didn’t attain symmetric MA-PIP2 interactions for two MA trimers in the immature MA complex (***Figure 3***). The lipid sorting behavior reported in the manuscript was verified using multiple trajectories.

### Molecular Dynamics Simulations and Analysis

All simulations used the CHARMM36m force field (71) and were performed in GROMACS 2019 MD software (72). Energy minimization was performed using the steepest descent algorithm until the maximum force was less than 1,000 kJ mol^−1^nm^−1^. Then, equilibration was performed with harmonic restraints (using a 1,000 kJ mol^−1^nm^−2^ spring constant) on each heavy atom throughout the protein for 1 ns in the constant NVT ensemble with a time step of 1 fs, followed by a 15 ns constant NVT MD run with a time step of 2 fs and a 15 ns MD run in the constant NPT ensemble with a time step of 2 fs. Next, each system was allowed to run for a total of 2000 ns and 3000 ns, respectively, for immature and mature MA complexes in the constant NPT ensemble, following the parallel cascade selection MD procedure, to allow Myr insertion to occur. During this phase, MA assembly structures were kept intact by applying harmonic restraints (1000 kJ mol^−1^nm^−2^) on Cα-backbone atoms of protein monomers, except for the first ten N-terminal residues including the Myr group. Next, 4 μs trajectories were generated for both the MA dimer of trimer complexes with all the six Myr groups inserted into the membrane. Previous restraints on the protein backbones were removed for production runs. For the immature MA complex, to mimic the effect of uncleaved Gag protein, we have applied small harmonic restraints (50 kJ mol^−1^nm^−2^) to the Cα-atom of ALA119 residues (C-terminal residue of helix 5). To further confer stability to the immature MA dimer of trimers assembly structure, harmonic restraints (100 kJ mol^−1^nm^−2^) were applied to the N-terminal residues (residues 4-8) of the four peripheral MA monomers (not at the trimer-trimer interface) in order to emulate the binding of immature MA lattice. No such restraints were applied to the mature MA complex. Among many replicas of the production runs, the attainment of specific protein-lipid interactions ensured the stability of the MA complexes.

Throughout this procedure, the temperature was kept constant at 310.15 K using the Nosé - Hoover thermostat with a 1.0 ps coupling constant (73,74), and the pressure was set at 1 bar and controlled using the Parinello-Rahman barostat semi-isotropically due to the presence of the membrane. The compressibility factor was set at 4.5 ×10^−5^ bar^−1^ with a coupling time constant of 5.0 ps (75,76). Van der Waals interactions were computed using a force-switching function between 1.0 and 1.2 nm, while long-range electrostatics were evaluated using Particle Mesh Ewald(PME) (77) with a cutoff of 1.2 nm, and hydrogen bonds were constrained using the LINCS algorithm (78).

In addition, we performed biased MD simulations to explore the conformational space of PS and PIP2 lipids at the trimer-trimer interface of immature and mature MA complexes [see *Supporting Information* for details]. Model fitting was done in ChimeraX. Protein-protein and protein-lipid interactions were visualized using VMD 1.9.3. (69). Analyses of the trajectories were performed using GROMACS 2019 (72) and VMD Tcl scripts(69). Lipid density maps were generated using the Python package Matplotlib (79).

### Biased molecular dynamics simulation details

To investigate the “extended lipid conformations” predicted by NMR experiments and cryoET density map, we have explored PIP2 and PS lipid conformational space employing steered MD (SMD) and restrained MD(rMD) simulations. PIP2 and PS lipids at the trimer-trimer interface for immature and mature MA assemblies were pulled out of the membrane from multiple initial configurations selected from unbiased MD trajectory. SMD simulations were carried out for 10 ns using constant velocity pulling (0.5 nm per ns) using a biasing force constant of 1000 kJ mol^−1^ nm^−2^. In these simulations, pulling the headgroup of lipids away from the membrane bilayer was not successful to sample extended lipid conformation. However, pulling the center of mass (COM) of each lipid molecule along the membrane normal, maintaining the MA-lipid(headgroup) interactions, sampled such extended lipid conformations for some selected PIP2 lipids. From each SMD trajectory, multiple configurations were selected depending on the distance between the COM of lipids and membrane bilayer, and restrained MD simulations (rMD) were performed for 150 ns using a harmonic potential of 1000 kJ mol^−1^ nm^−2^ along the reaction coordinate. Total ∼20 μs of SMD and rMD simulations were carried out and finally, the stability of extended lipid conformations obtained for mature and immature MA assemblies was assessed by performing unbiased MD simulations (1 μs).

## Author Contributions

P.B., J.A.G.B, and G.A.V. designed the research. P.B. prepared simulation models, designed and performed simulations, analyzed the data, prepared the figures, and wrote the manuscript. K.Q. contributed to the discussions and manuscript preparation. J.A.G.B and G.A.V. supervised the research, writing process, and contributed to the discussions to finalize the manuscript.

## Acknowledgments

This research was supported by the National Institute of Allergy and Infectious Diseases (NIAID) of the National Institutes of Health (NIH grant R01AI178550) to G.A.V. and J.A.G.B. and the Max Planck Society to J.A.G.B. The authors acknowledge computational resources provided by the University of Chicago Research Computing Center (RCC), the Frontera supercomputer at the Texas Advanced Computer Center funded by the National Science Foundation (OAC-1818253), the Stampede2 supercomputer at the Texas Advanced Computing Center (TACC), the Bridges2 supercomputer at the Pittsburgh Supercomputing Center (PSC) through allocation MCA94P017 with resources provided by the Extreme Science and Engineering Discovery Environment (XSEDE) supported by NSF grant ACI-1548562 and the NIH-funded Beagle-3 computer (NIH award 1S10OD028655-01).

