## Supplementary Materials for "Molecular dynamics simulations of HIV-1 matrix-membrane interactions at different stages of viral maturation"

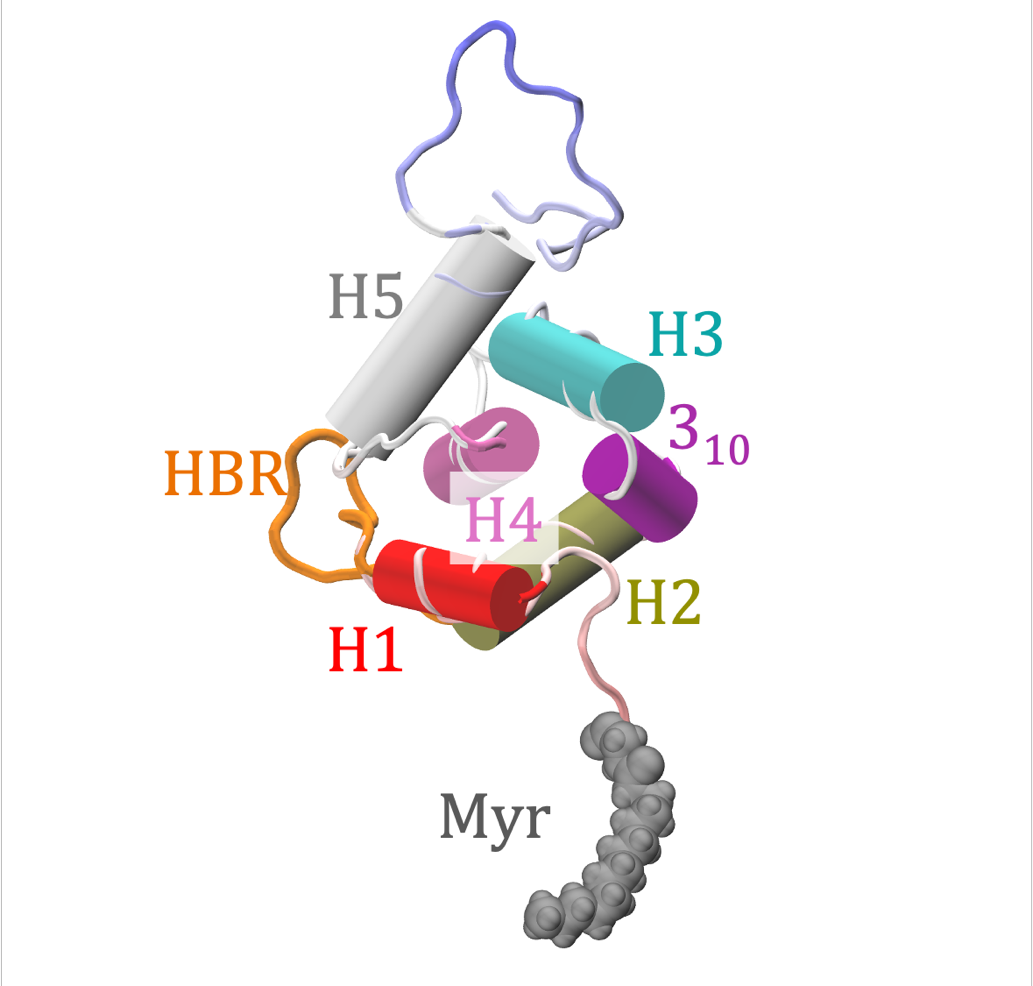

**Figure S 1: Side-view of a membrane-bound MA monomer with Myr group in exposed conformation.** Key regions are labeled: H1, H2, 3_10_, H3, H4, and H5 helices (in order of appearance in the protein sequence). HBR(highly basic region), shown as an orange flexible loop, connects H1 and H2 helices. The Myr group, shown in gray, is covalently attached to the N-terminal glycine residue. In our simulated model, residue 1 is composed of Myr group and GLY residue. Therefore, residue­_n_ of the experimental PDB structure corresponds to residue_n-1_ of our simulated models.

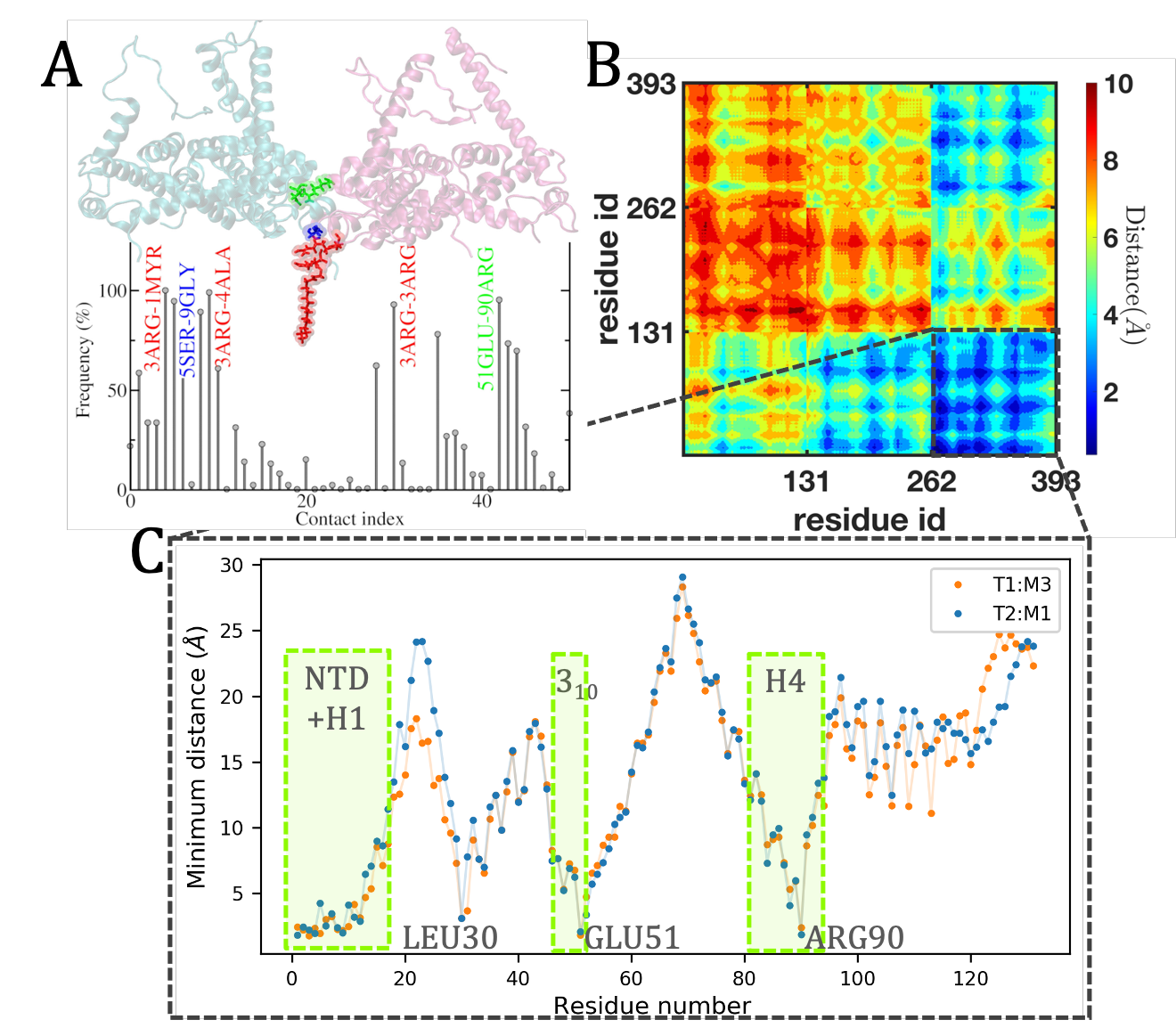

**Figure S 2: Trimer-trimer interactions (TTIs) in the immature MA assembly.** A. Frequency of contacts among residues of two trimers (cut-off distance 2.5 Å). Pairs of residues with a contact frequency of more than 95% are highlighted. B. Distance map of two trimers. Residues 1-131, 132-262, and 263-393 constitute monomer1, monomer2, and monomer3 respectively. Monomer3 of trimer1 (T1:M3) is in contact with monomer1 of trimer2 (T2:M1) in our simulated model. C. Minimum distances of the other trimer from the residues of T1:M3 and T2:M1. N-terminal residues, H1, 3_10,_ and H4 residues of both the monomers stabilize the trimer-trimer interface.

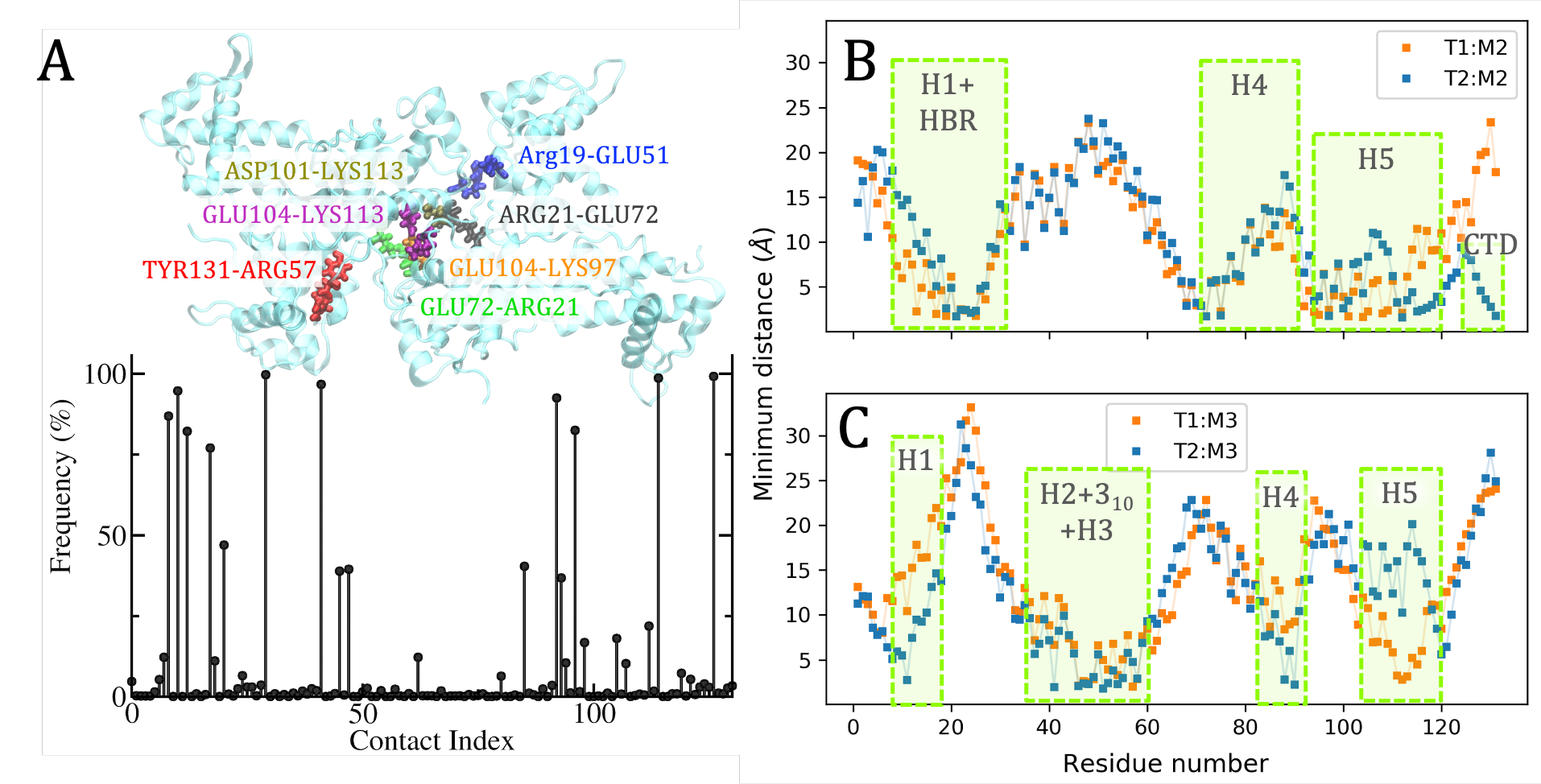

**Figure S 3: Trimer-trimer interactions (TTIs) in mature MA assembly.** A. Frequency of contacts among residues of two trimers (cut-off distance 2.0 Å). Pairs of residues with a contact frequency of more than 95% are highlighted. In this assembly, four monomers from both the trimers(T1&T2) are in contact with each other. Minimum distances of the other trimer from the residues of each of these monomers are computed. B. Monomer2 of both the trimers (T1:M2, T2:M2) participate in the trimer-trimer interactions mainly with the HBR domain, H4, H5. Unlike immature MA, in this simulated system residues of H5 helices participate in the trimer-trimer interactions. C. Monomer3 of both the trimers (T1:M3, T2:M3) interact with the other trimer via residues of H2, 3_10,_ and H3 helices.

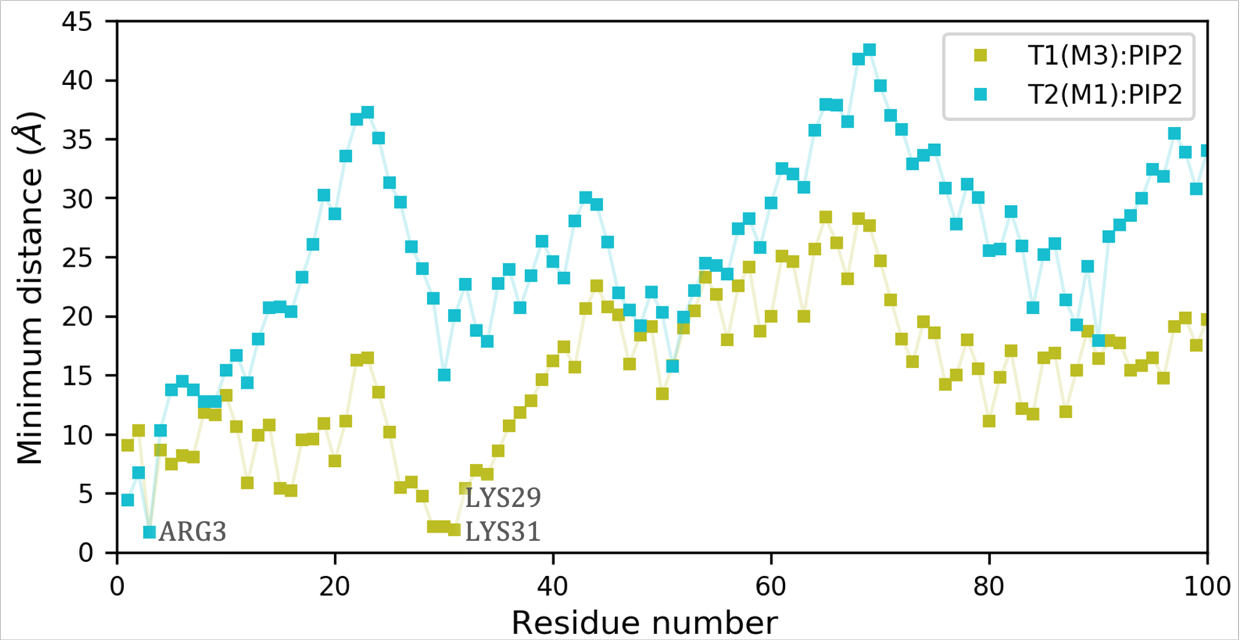

**Figure S 4:**  **MA-PIP2 interactions at an alternate PIP2 binding site of the immature MA assembly.** Minimum distances of the headgroup of PIP2 lipid at the binding site ‘a’ (shown in Figure 3) from the residues of two MA monomers at the interface, averaged over the trajectories. ARG3 of both the monomers at the trimer-trimer interface (T1:M3 and T2:M1) binds to the headgroup of PIP2. Apart from those, positively charged residues like LYS29, and LYS31 of T1:M3 interact with the negatively charged inositol headgroup of that PIP2. This simulated MA-PIP2 binding agrees with NMR experimental data(35).

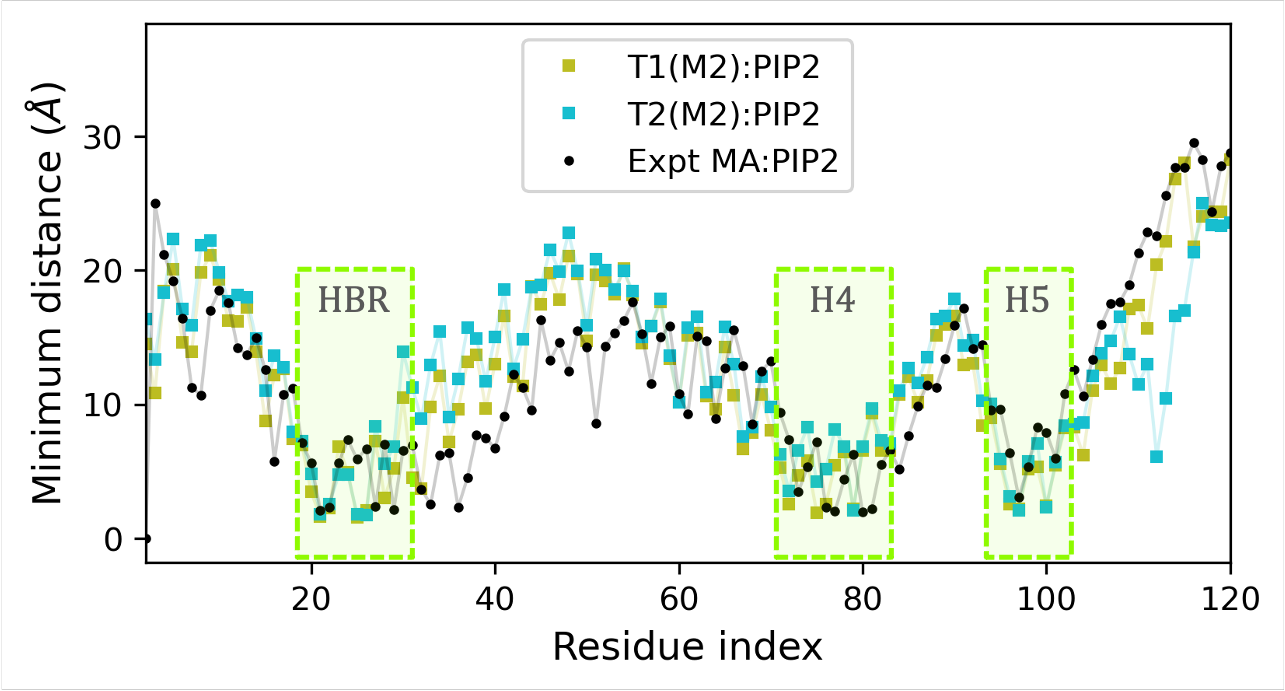

**Figure S 5: MA-PIP2 interactions at the trimer-trimer interface of mature MA assembly.** The minimum distance of two PIP2 molecules in an extended lipid conformation at the binding site ‘a’ (shown in Figure 4), averaged over the trajectories. PIP2 binding with monomer2 of both the trimers (T1:M2, T2:M2) agrees well with experimentally predicted PIP2 binding (PDB ID: 2H3V)(29).

**Figure S6:** Radial distribution function (rdf) of headgroup phosphorus(P) atoms of PS lipids around the headgroup P atoms of PIP2 lipids. The RDF profile shown in black is computed for 5 specific PIP2 lipids which are bound to MA monomers at the trimer-trimer interface (TTI) of the mature MA assembly. The RDF profile shown in pink is computed for all other PIP2 lipids in the system except for those 5 PIP2 lipids at TTI. RDF profiles are calculated using GROMACS tool g_rdf.
